# scFv intrabody targeting wildtype TDP-43 presents protective effects in a cellular model of TDP-43 proteinopathy

**DOI:** 10.1101/2025.03.19.644089

**Authors:** Yara Al Ojaimi, Rudolf Hergesheimer, Anna A. Chami, Hugo Alarcan, Johanna Augros, Audrey Dangoumau, Shanez Haouari, Jérôme Bourgeais, Antoine Lefevre, Samira Osman, Patrick Emond, Patrick Vourc’h, Christian R. Andres, Philippe Corcia, Olivier Herault, Pierre Martineau, Débora Lanznaster, Hélène Blasco

## Abstract

TDP-43 proteinopathies are a set of neurological disorders characterized by the abnormal accumulation and mislocalization of TDP-43 in the cytoplasm, leading to the disruption of the normal function of the protein. In most of the cases, it is the wildtype (wt) form of the protein that is involved. An untargeted high-throughput screen of a single-chain variable fragment (scFv) library was performed using phage display against human full-length wt TDP-43. Two scFvs (B1 and D7) were retained following cellular expression (then termed intrabodies) and colocalization with cytoplasmic TDP-43 *in vitro.* We generated a 3D structure of full length wt TDP-43 *in silico*, and used it for epitope mapping. In a cellular model of TDP-43 proteinopathy, D7 enhanced the proteasomal degradation of the insoluble 35-kDa C-terminal fragment TDP-43 and reversed some TDP-43-induced metabolomic alterations, particularly relating to the lipid metabolism. Our findings offer a new scFv intrabody that bind to human wtTDP-43 and modify cellular pathways associated with TDP-43 proteinopathies.

## 1. Introduction

TDP-43 is a DNA/RNA-binding protein encoded by the TARDBP gene, and is essential for cell homeostasis(1,2). Under physiological conditions, TDP-43 is mostly localized in the nucleus and shuttles between the nucleus and cytoplasm. TDP-43 was mainly linked to amyotrophic lateral sclerosis (ALS), where 97% of patients present with cytoplasmic aggregates of the protein, regardless of their mutational status(2). However, the protein was later found to be dysregulated in several other neurodegenerative diseases like frontotemporal dementia (FTD)(3), Alzheimer’s(4), Parkinson’s(5), and Huntington’s disease(6), and limbic-predominant age-related TDP-43 encephalopathy (LATE)(7). These diseases became therefore collectively known as TDP-43 proteinopathies. In these diseases, it is TDP-43 in its wild-type (wt) form that is found mislocalized and aggregated in the cytoplasm.

TDP-43 aggregates mainly include full-length and 35-kDa and 25-kDa C-terminal fragments (CTF) of TDP-43(3,8). The C-terminal fragments are a result of proteolytic cleavage or alternative splicing and have been widely implicated in toxic gain- or loss-of function mechanisms(9). Dysregulation of TDP-43 was found to induce metabolic alterations, particularly in pathways involving energy metabolism, lipids, and neurotransmission(10,11). Moreover, there seems to be a bidirectional interplay between the function of the degradation pathways of the cell, particularly the ubiquitin-proteasome system and TDP-43 proteinopathy, suggesting a key role of these pathways in TDP-43 dysfunction(12,13). However, targeting TDP-43 pathology is complicated by the plethora of cellular pathways affected by the protein and its role in maintaining cellular homeostasis (14). For instance, knockdown of TDP-43 by siRNA was found to be associated with impaired axonal transcriptome(15), increased inflammation(16,17), and dysregulation of the expression of important proteins implicated in cell homeostasis(18,19). Similarly, antisense oligonucleotides (ASO) that downregulate TDP-43 in mice resulted in synaptic dysfunction and increased neuronal vulnerability(1).

Researchers are therefore trying to develop new strategies such as single-chain variable fragments (scFvs) to better regulate this protein is TDP-43 proteinopathies.

scFvs are generated by fusing the variable V_H_ and V_L_ regions of an IgG antibody with a linker peptide, resulting in an antibody fragment of 25-35 kDa that lacks the constant regions of the 150-kDa IgG yet still retains antigen-binding specificity. Due to their small size, scFvs haven been used as intracellular antibodies, called intrabodies. In recent years, important developments have been made in the use of scFvs for passive immunization targeting proteinopathies in many neurodegenerative diseases(20–25).

Our aim for this project was to apply an innovative, untargeted high-throughput strategy based on a scFv screening method to target full length wtTDP-43, and to evaluate diverse cellular variables to elucidate their effects. We used phage display to screen an scFv library against recombinant human full-length wt TDP-43. Since no 3D structure of full length TDP-43 was available, one was generated *in silico* externally, which we used to identify pathologically relevant epitopes. The retained scFv was investigated for its cellular localization, its impact on cell viability and the expression and solubility of TDP-43, as well as its effect on other subtle cellular alterations that might occur earlier in TDP-43 proteinopathies, such as metabolic alterations.

## 2. Materials and Methods

### 2.1 Plasmids and cloning

We had previously cloned a full-length human wtTDP-43 cDNA(26) into the pcDNA3.3-TOPO TA vector (Invitrogen). Starting from this clone, we added a 6xHis tag to the 3′ end by performing PCR on the plasmid using the following primers (Eurogentec): forward (5′-TCTGAATATATTCGGGTAACCGAAG-3′) and reverse (5′-CTAGTGATGGTGATGGTGATGAGAACCCCCCATTCCCCAGCCAGAAGACTTAG–3′). The resulting amplicon was then integrated into the Vivid Colors pcDNA 6.2/C-EmGFP-GW/TOPO Mammalian Expression Vector (Invitrogen), which added GFP to the 5′ end and yielded GFP-wtTDP-43-6xHis to permit mammalian cell expression and downstream purification for phage display. To clone the sequence for the control antigen for phage display, GFP-6xHis, a similar approach was taken by first performing PCR on the plasmid containing GFP-wtTDP-43-6xHis using the following primers (Eurogentec): forward (5′– GCCACCATGGTGAGCAAGGGCGAGGA–3′) and reverse (5′– CTAGTGGTGATGGTGATGATGAGAACCCCCCTTGTACAGCTCGTCCATGCC–3′). Then, the GFP-6xHis sequence was integrated into an empty pcDNA3.3-TOPO plasmid.

As for scFv cloning, using the NcoI and NotI restriction sites surrounding the 5′ and 3′ ends of the scFv cDNA sequence, respectively, the pHEN-HuscI plasmid was digested to completion. The liberated scFv sequence was then integrated into previously digested empty pET-23NN vector using T4 DNA Ligase (Thermo Fisher Scientific). This vector added a c-myc and 6xHis tag at the 3′ end of the clone, which permitted expression and purification from BL21(DE3)pLysS bacteria (New England BioLabs). From the integrated pET-23NN vector, we cloned the scFv sequence with the 3′ tags into pcDNA3.3-TOPO plasmid to permit mammalian cell expression according to the manufacturer’s instructions (Thermo Fisher Scientific). For the PCR step, the following primers were used (Eurogentec): forward (5′– GCCACCATGGCCGAGGTGCAGCTG–3′) and reverse (5′-CTAATGGTGATGATGGTGATGTGCGG–3′).

### 2.2 Mammalian Cell Culture

HEK-293T human cells (ATCC) were the main cell line of choice due to its robust transfection efficiency and common application in studies on TDP-43 proteinopathy. We maintained cells in Dulbecco’s Modified Eagle’s Medium (DMEM) supplemented with 5% (v/v) fetal bovine serum (FBS) at 37 °C in an incubator maintaining an atmosphere of 5% CO_2_. Cells were split once they reached 90% confluency.

### 2.3 Purification of antigens from HEK-293T cells

The recombinant proteins GFP-6xHis and GFP-wtTDP-43-6xHis were overexpressed for 48h in HEK293T cells plated at a density of 0.44 × 10^6^ cells/T-150 flask. At 70% confluency, the cells were transfected with 60 µg of expression plasmid mixed with Lipofectamine-2000 in a ratio of 1:2 (Invitrogen), following the manufacturer’s protocol. The cells were harvested and centrifuged at 500 × g for 10 min at 4° C then resuspended in cold denaturing lysis buffer (500 mM NaCl, PBS pH 8.0, 10% glycerol, 0.1% Brij-35, 6 M urea, 1 mM DTT) supplemented with 1X Halt Protease Inhibitor Cocktail (Thermo Fisher Scientific). Four freeze-thaw cycles using liquid nitrogen and a 37° C water bath were applied to the cells, then Pierce Universal Nuclease (250 U/µL, Invitrogen) and 10 mM MgCl_2_ were added to the lysate. After 15 min of agitation at 4° C, the lysate was centrifuged at 15,000 × g for 20 min at 4° C. The supernatant was passed through an equilibrated Ni-NTA resin column (Thermo Fisher Scientific). The column was then washed with cold lysis buffer supplemented with 25 mM imidazole. Finally, the recombinant protein was eluted by the addition of cold lysis buffer supplemented with 250 mM imidazole. The fractions containing recombinant protein were identified by SDS-PAGE and pooled.

The pooled eluate was transferred to dialysis tubing (CelluSepH1, Membrane Filtration Products) with a molecular-weight cutoff (MWCO) of 15 kDa and incubated in 500 mL of dialysis buffer (50 mM Tris-HCl pH 7.4, 10% glycerol, 150 mM NaCl, 0.1 mM EDTA, 2 mM β-mercaptoethanol) at 4° C for 16 h to permit the refolding of the purified protein. The buffer was then replaced with 500 mL of fresh buffer for another incubation for 2 h at 4° C. The dialyzed solution containing refolded protein was concentrated using an Amicon Ultra tube (Merck Millipore) with a MWCO of 10 kDa at 4° C. The concentration of the purified refolded protein was determined by Bradford assay (Thermo Fisher Scientific) following the manufacturer’s protocol.

To characterize the size of the purified recombinant proteins, a sample of 100 µL was injected into an FPLC AKTA Purifier10 (GE Healthcare) with a 25-mL Superdex 200 Increase 10/300 GL column (GE Healthcare) equilibrated in dialysis buffer at 4° C; 500-µL fractions were collected at a flow rate of 0.5 mL/min. The identity of the purified recombinant protein was confirmed by western blot using a 6xHis tag antibody (HRP-66005, Proteintech).

### 2.4 HuscI phage library and control scFv

The HuscI (Human scFv Intrabody Library) library provided by Dr. Pierre Martineau from the Montpellier Cancer Research Institute(27–29) is a human synthetic library with a diversity of 3 × 10^9^ functional clones. The library is cloned in vector pHEN1 (pHEN-HuscI), resulting in M13 phage clones expressing single-chain variable fragments (scFvs) at the N-terminal extremity of pIII protein. Each scFv consists of a variable heavy (VH) and light (VL) chain connected by a linker sequence (GGGGSGGGGSGGGGS). The molecular weight of the scFvs ranges from 25 to 30 kDa. The scFv employed as the control in this work is the anti-β-galactosidase scFv, termed 13R4, described by Martineau and his colleagues(27).

### 2.5 Phage display and polyclonal selection

Before panning on GFP-wtTDP-43-6xHis (GTH), the HuscI scFv library was depleted on anti-GFP and GFP-His protein. Three wells of a Nunc Maxisorp 96-well plate (Thermo Fisher Scientific) were coated with 10 µg/mL of anti-GFP antibody (Roche, 11814460001) diluted in PBS, for 24 h at 4 °C. After 5 washes in PBS-Tween 0.1% (PBST) and one wash in PBS, the wells were saturated in 4% powdered milk in PBS (MPBS) for 2 h at RT. After washing, the first well was incubated with 50 µL of MPBS and 50 µL of the HuscI phage library. In parallel, the second well was incubated with 10 µg/mL of purified GFP-6xHis diluted in 100 µL of PBST. After 2 h, the supernatant from the first well was transferred to the second well for another 2-h incubation. During this time, the third well was incubated with 5 µg/mL of purified GFP-wtTDP-43-6xHis diluted in 100 µL of PBST. After 2 h, the supernatant from the second well was transferred to the third well to begin the first round of selection.

Following 2 h of incubation in the third well at RT, the supernatant was discarded and the well was washed 20 times with PBST and 3 times with PBS. Next, 100 µL of elution buffer (50 mM Tris pH = 8.0, 1 mM CaCl_2_) containing 125 µg/mL of trypsin (TPCK-treated, Sigma-Aldrich) was added to the well for 15 min at RT. The supernatant was transferred to 200 µL of 2xYT medium (Sigma-Aldrich) containing TG1 *E. Coli* at an optical density (OD) of 0.5. After 1 h at 37 °C with shaking, the bacteria were centrifuged at 3000 × g for 10 min at RT and resuspended in 500 µL of 2xYT/2% glucose/100 µg mL^−1^ ampicillin. The bacteria were allowed to grow at 30 °C with shaking for 16 h, then 20 µL of the culture were added to 1 mL of fresh 2xYT/2% glucose/100 µg mL^−1^ ampicillin and shaken at 37 °C. After reaching an OD of 0.5, the culture was infected with 20 µL of KM13 helper phage(30) comprising approximately 10^11^ TU/mL. Following shaking at 37 °C for 1 h, the bacteria were centrifuged at 3000 × g for 10 min at RT and resuspended in 1 mL of 2xYT with 100 µg/mL ampicillin and 25 µg/mL kanamycin. The culture was shaken at 30 °C for 16 h, then 500 µL of this culture was centrifuged at 10,000 × g for 10 min at 4 °C. Next, 400 µL of the supernatant containing new phage was added to 100 µL of 20% polyethylene glycol-8000 (PEG) and sterile 2.5 M NaCl at 4°C to precipitate the phage. After 15 min on ice, the solution was centrifuged at 13,000 rpm for 20 min at 4 °C. The pellet was resuspended in 200 µL of 15% sterile glycerol diluted in PBS. The phage titer was calculated using the following formula:

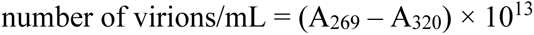

Next, 50 µL of the new phage stock containing approximately 10^11^ to 10^12^ phage was added to 50 µL of MPBS in a new well coated with 5 µg/mL of GFP-wtTDP-43-6xHis to undergo the second round of selection. In total, 4 rounds of selection were performed.

### 2.6 Phage display: monoclonal selection

A flowchart of the phage display strategy for identifying the hits against wtTDP-43 is represented in **Fig.S1**. HB2151 cells (a strain allowing the expression of soluble unfused scFv, provided by Dr. Pierre Martineau’s group) at an optical density of 0.5 were infected with 10^6^ phage from polyclonal selection round 4. After 30 min at 37 °C without shaking, infected bacteria were plated on LB/agar with 1% glucose and 100 µg/mL ampicillin and incubated at 37 °C for 16 h. Ninety-six isolated HB2151 colonies were transferred to wells of a 96-well deep-well plate (Thermo Fisher Scientific) containing 300 µL of 2xYT, 100 µg/mL ampicillin, and 1% glucose. The plate was incubated at 37 °C with agitation for 16 h, then 2–5 µL of culture from each well was transferred to a new flat-bottom plate (Thermo Fisher Scientific) containing 200 µL of 2xYT, 100 µg/mL ampicillin, and 0.1% glucose in each well. At an optical density of 0.6, 25 µL of fresh medium supplemented with 9 mM IPTG (Sigma-Aldrich) was added to each well for induction. Induction lasted 24 h at 30 °C with shaking.

The plate was centrifuged at 1800 × g for 10 min. The supernatant contained soluble scFv with a C-terminal myc tag; 50 µL of each supernatant were added to 50 µL of 4% MPBS in each well of a 96-well plate pre-coated with 1.0 µg/mL of GTH and GH antigen (2 plates total). After a 1-h incubation at RT, the wells were washed 3 times in PBS, and 100 µL of anti-c-Myc antibody (9E10, Thermo Fisher Scientific) diluted 2000-fold in saturation solution was added to each well and incubated for 1 h at RT. The wells were washed 3 times with PBST and 1 time with PBS, and 100 µL of TMB-ELISA Substrate Solution (Thermo Fisher Scientific) was then added for an incubation of 5–15 min at RT in the dark. The reaction was halted with 50 µL of 1 M H_2_SO_4_. Absorbance was measured at 450 nm using an iMark Microplate Absorbance Reader (Bio-Rad).

For the alternative monoclonal selection, the glycerol stock of TG1 infected with phage from round 4 of selection was incubated in 2xYT/2% glucose/100 µg/mL ampicillin for overnight shaking at 37 °C. A small volume of the culture was then plated on LB agar with 100 µg/mL ampicillin. After 16 h at 37 °C, 95 colonies were transferred to each well of a deep-well 96-well plate containing 300 µL of 2xYT/1% glucose/100 µg mL^−1^ ampicillin. Following 16 h of shaking at 37 °C, 5 µL of culture from each well was transferred to another flat-bottom 96-well plate with 200 µL of 2xYT/2% glucose/100 µg mL^−1^ ampicillin. At an OD of 0.5, the cultures were infected with KM13 helper phage, as described previously.

The cultures were then centrifuged at 3000 × g for 10 min and resuspended in fresh medium supplemented with 25 µg/mL kanamycin. After 16 h at 30 °C with shaking, the cultures were centrifuged again, and 50 µL of supernatant was added to 50 µL of MPBS and incubated in a 96-well plate pre-coated with 1.0 µg/mL GFP-6xHis and another plate pre-coated with 1.0 µg/mL GFP-wtTDP-43-6xHis. Following 1 h at RT, the wells were washed 3 times with PBST and 1 time with PBS, and 100 µL of 4% MPBS containing HRP-conjugated anti-M13 bacteriophage antibody (Sino Biological) at 1/2000 dilution was added to each well for an incubation at RT for 1 h. The same washing procedure was applied before adding 100 µL of TMB-ELISA Substrate Solution (Thermo Fisher Scientific) for an incubation of 5–15 min at RT in the dark. The reaction was halted with 50 µL of 1 M H_2_SO_4_. Absorbance was measured at 450 nm.

### 2.7 In silico epitope mapping

The potential binding sites of the scFv of interest were predicted *in silico* for wtTDP-43 by MAbSilico (Nouzilly, France). Given that no 3D structure of full-length TDP-43 was found, it was generated by combining the resolved structures of the separate domains of TDP-43 and homologous proteins available on the RCSB PDB Protein Data Bank. Residues 1–88 were modeled by region 1–102 of TDP-43 (PDB file 5MRG)(31). Residues 89–102 were modeled by the structure of nuclear ribonucleoprotein L (PDB file 2MQN)(32). Residues 103–177 and 191–258 were modeled by both RRM domains of TDP-43 (PDB file 4BS2)(33). Residues 259– 306 were modeled by U1 small nuclear ribonucleoprotein 70 (PDB file 5UZ5)(34). Residues 307–346 were modeled by the hydrophobic helix, prion-like domain of TDP-43 (PDB file 2N2C). Finally, residues 347–413 were modeled by human splicing factor (PDB file 4WIJ)(35). Briefly, epitope mapping was based on a docking method(36). The probability of an interaction for each residue involved in the top 30 conformations of the scFv-TDP-43 complex was also evaluated.

### 2.8 Overexpression of recombinant proteins in mammalian cells

Six-well plates (CytoOne) were each seeded with 0.3 × 10^6^ cells in DMEM/5% FBS/pyruvate/1% non-essential amino acids. When the cells reached 60% confluency, they were transfected for 48h (Polyplus transfection) using jetPEI transfection reagent (Polyplus transfection) following the manufacturer’s protocol. Regarding transfections, a total of 4.5 µg of plasmids were always used. For the TDP-43 condition, 3.0 µg of wt-TDP-43-expressing plasmid with 1.5 µg of empty plasmid was used. For the intrabodies alone condition, 1.5 µg of scFv-expressing plasmids and 3 µg of empty plasmid were used. With respect to TDP-43/scFv co-transfections, 3.0 µg of wtTDP-43 was mixed with 1.5 µg of scFv-expressing plasmid. Unless noted otherwise, all TDP-43 transfections included the untagged, full-length, wild-type form.

### 2.9 Immunofluorescence

For immunofluorescence experiments, an 8-well Nunc Lab-Tek II plate (No. 1.5 borosilicate glass, Thermo Fisher Scientific) that was pre-coated with 50 µg/mL of poly-D-lysine (Sigma-Aldrich) was seeded with 0.02 × 10^6^ HEK293T cells. After 24 h, transfection was performed as described previously with 0.2 µg of TDP-43 plasmid or empty plasmid and 0.1 µg of scFv plasmid. Following 48 h of overexpression, the cells were fixed in 4% paraformaldehyde and 2% sucrose for 20 min at RT. The cells were washed 3 times in PBS. The fixed cells were permeabilized using 0.2% Triton-X 100 diluted in PBS for 15 min at RT. Then they were saturated in permeabilization buffer supplemented with 10% donkey serum (ab7475, Abcam) for 1 h at RT. Next, 1000-fold diluted rabbit anti-TDP-43 primary antibody (12892-1-AP, Proteintech), and 1000-fold diluted mouse anti-6xHis primary antibody (66005-1-Ig, Proteintech) were added to the cells for an incubation of 2 h at RT. Cells were washed 5 times the incubated wtih 500-fold diluted goat anti-rabbit Alexa Fluor 488 (A-11034, Thermo Fisher Scientific) and 500-fold diluted donkey anti-mouse Alexa Fluor 594 (A-11005, Thermo Fisher Scientific) for 1 h at RT away from light. Following 4 washes, the slides were mounted using Vectashield Antifade Mounting Medium with DAPI (H-1200-10, Vector Laboratories). then visualized with a SP8 confocal microscope (Leica Microsystems) using a magnification of 60X. A table of all the antibodies used in this study is presented in **Table S1**.

### 2.10 Cell viability

Following 48 h of overexpression, the culture medium was withdrawn and replaced with 0.5 mg/mL MTT (3-(4,5-dimethylthiazol-2-yl)-2,5-diphenyltetrazolium bromide, Sigma-Aldrich) diluted in HBSS (Gibco). After 30 min of incubation at 37 °C with 5% CO_2_, the medium was discarded and replaced with 500 µL of DMSO (Sigma-Aldrich), in which the formazan crystals were dissolved. The absorbance was recorded at 570 nm.

### 2.11 Purification of scFv from BL21(DE3)pLysS

A culture for a given clone was started in LB containing 1% glucose and 100 µg/mL ampicillin and incubated for 16 h at 37 °C with shaking. At an OD of 0.6, 12 mL of culture was added to an auto-inducible medium consisting of fresh LB containing 100 µg/mL ampicillin, 2 mM MgSO_4_, 20mM 50x5052 (2.7 M glycerol, 0.14 M glucose, 0.28 M α-lactose monohydrate), and 20mM of solution 50xM (1.3 M Na_2_HPO_4,_ 1.3 M KH_2_PO_4_, 2.5 M NH_4_Cl, 0.25 M Na_2_SO_4_).

The culture was incubated at 37 °C with shaking for 3 h, then moved to 22 °C for 16 h. The bacteria were pelleted by centrifugation at 5000 × g for 10 min at 4 °C and resuspended in lysis buffer (PBS pH 8.0, 1% Brij-35, 1.0 mM EDTA, 10 mM MgCl_2_, 1.0 mM DTT). To this solution were added 1% Halt Protease Inhibitor Cocktail (Thermo Fisher Scientific), Pierce Universal Nuclease (Thermo Fisher Scientific), and 0.1 M of lysozyme. After 15 min of agitation at 4 °C, the bacteria were lysed by sonication. The lysate was centrifuged at 15,000 × g for 20 min at 4 °C. The supernatant was passed through a of Ni-NTA resin column (Thermo Fisher Scientific) equilibrated in lysis buffer with 10 mM imidazole. The column was then washed with cold lysis buffer supplemented with 25 mM imidazole. The scFv was eluted using cold lysis buffer supplemented with 250 mM imidazole. Fractions containing the scFv were identified by SDS-PAGE and pooled, then dialyzed in MilliQ water. The dialyzed product was quantified by Bradford assay and concentrated to 0.5–1.0 mg/mL if necessary. The final product was stored at 4 °C. The identity of the scFv was confirmed by western blot.

### 2.12 Enzyme-linked immunosorbent assay (ELISA) on purified scFv

Wells of a Nunc MaxiSorp 96-well plate (Thermo Fisher Scientific) were coated overnight at 4 °C with commercial human wtTDP-43 (Abcam, ab224788) diluted to 1 µg/mL in PBS. Wells were washed 3 times in 0.2% PBST, and blocked using PBS Tween-20(0.02%) and 5% fat-free milk for 2 h at RT. The wells were washed 3 times then incubated with purified scFv diluted to 1, 5, or 10 µg/mL for 1 h at RT. Following 3 washes, HRP-conjugated mouse anti-6xHis antibody (66005, Proteintech) diluted 10,000-fold in PBS-T(0.02%) was then added to the wells and incubated for 1 h at RT, after which the wells were washed 5 times. Next, TMB-ELISA Substrate Solution (Thermo Fisher Scientific) was added for an incubation of 10– 15 min at RT in the dark. The reaction was halted with 2 M H_2_SO_4_. Absorbance was measured at 450 nm using an iMark Microplate Absorbance Reader (Bio-Rad).

### 2.13 TDP-43 solubility

Following 48 h of overexpression in HEK293T, the cells were lysed using ice cold lysis buffer (PBS pH 8.0, 1% Triton-X 100, 10 mM MgCl_2_, 1 mM DTT) supplemented with 1X Halt Protease Inhibitor Cocktail (Thermo Fisher Scientific) and Pierce Universal Nuclease (Thermo Fisher Scientific). After a 30 min incubation at 4 °C, the lysates were centrifuged at 13,000 × g for 20 min at 4 °C. The supernatant (soluble fraction) was transferred to a new tube, while the pellet (insoluble fraction) was resuspended in the same volume as the supernatant of ice cold lysis buffer supplemented with 6 M urea. The protein concentrations of the crude extracts of the lysates were measured by the detergent-compatible Pierce 660nm Protein Assay reagent (Thermo Fisher Scientific). For western blot analysis, 10 µg of crude extract along with equivalent volumes of the insoluble fractions, were run on an SDS-PAGE gel, transferred to a PVDF membrane, and blocked in TBS-Tween20 (0.1%). TDP-43 was stained overnight at 4 °C by a rabbit primary antibody targeted against the C-terminus (12892-1-AP, Proteintech) diluted 5000-fold. Then, a 1-h incubation at RT was performed with HRP-conjugated goat anti-rabbit secondary antibody (W401B, Promega) diluted 10,000-fold. The 6xHis tag of the scFv was stained for 1 h at RT with the HRP-conjugated mouse anti-6xHis antibody (66005-1-Ig, Proteintech) diluted 100,000-fold.

### 2.14 TDP-43 degradation

Following wtTDP-43 and/or scFv overexpression in HEK293T cells, the medium was replaced with OptiMEM (Gibco, Thermo Fisher Scientific) and the cells were treated for 6 hours with 10 µg/mL cycloheximide (Sigma-Aldrich) with either 300 nM bafilomycin A1 (Merck, Sigma-Aldrich) or 0.5 µM MG-132 (Sigma-Aldrich). Protein extraction and western blots were performed as described previously.

### 2.15 Metabolomics analysis

After 48 h of overexpression in HEK293T, the cells were harvested and washed twice in ice-cold PBS. The dry cell pellet was frozen at −80 °C until the day of analysis. Cells were thawn on ice and resuspended in 150 µL of MeOH:H_2_O (1:1) followed by centrifugation for 5 min at 4 °C. Liquid chromatography in tandem with high-performance mass spectrometry (LC-HPMS) was performed as previously described by our group(26,37).

### 2.16 Data analyses

Unless otherwise stated, results are shown as mean ± standard deviation (SD). When relevant, the Mann–Whitney non-parametric t-test or an ANOVA test was performed using Prism v.7.0 (GraphPad Software, San Diego, CA, USA). Results were considered significant with *p* < 0.05.

For metabolomics analysis, each overexpression condition was normalized to the empty vector condition and the mean of the normalization ratios was calculated. We selected the 10% of the metabolites that increased the most and the 10% of the metabolites that decreased the most. A Venn diagram allowed us to identify the common and specific metabolites impacted by the scFv in comparison with those in naïve cells. We also normalized within each series to the “TDP-43 condition” and conducted the same process to observe the metabolites impacted by the scFv on TDP-43 overexpression. Moreover, we built the Venn diagrams between the increased and the decreased metabolites on scFv-TDP-43 versus scFv-empty vector. Finally, among the metabolites impacted by TDP-43 overexpression alone, we selected the ones also impacted by the scFv.

## 3. Results

### 3.1 Identification of four anti–TDP-43 clones

The purification of recombinant GFP-TDP43-6xHis antigen for phage display yielded highly pure monomer **(Fig.S2)**. Following four rounds of bio-panning, we observed the highest enrichment of bound phage at round 4 (**Fig.S3.A**). Therefore, round 4 phage were selected for the monoclonal screen phase. ELISA of 96 retained phage from round 4 revealed 9 clones showing strong positive signals for TDP-43 and not for the control GFP-6xHis antigen, suggesting specific binding to TDP-43. Fifty four clones displayed positive signals for GFP, and 33 clones showing no signal (**Fig.S3.B,C**). Sanger sequencing of the V_H_ and V_L_ domains from the 9 phages revealed 4 distinct scFv clones, which we named B1, A2, B6, and D7.

### 3.2 Generation of a 3D model of TDP-43 and prediction of scFv binding sites

Given that no 3D structure of the full-length TDP-43 protein was found, a homology model was built using 6 different structures **(Fig. 1A)**. Difficulties were encountered with the relative orientations of the different domains **(Fig. 1B)**. The relative orientation of the N-terminal and RRM1 domains and structure of the hinge 1 region were taken from the resolved structure of hnRNP L (PDB ID: 2MQN), which consists of two RRM domains, the first having homologies with the N-terminal domain of TDP-43. The relative orientation of the RRM2 and prion-like domains and the structure of the hinge 2 region were taken from *S. cerevisiae* U1 snRNP (PDB ID: 5UZ5), which consists of one RRM domain, a hinge, a prion-like domain, and a C-terminal helix. The C-terminal helix was modeled from human splicing factor (PDB ID: 4WIJ), which consists of one RRM domain, a hinge, a prion-like domain, and a C-terminal helix. Since the homologies in these regions were stronger with 5UZ5, it was used to model their orientations. However, the C-terminal helix of 4WIJ is much longer and was therefore used to model the part of this helix that is absent from 5UZ5.

**Figure 1:**
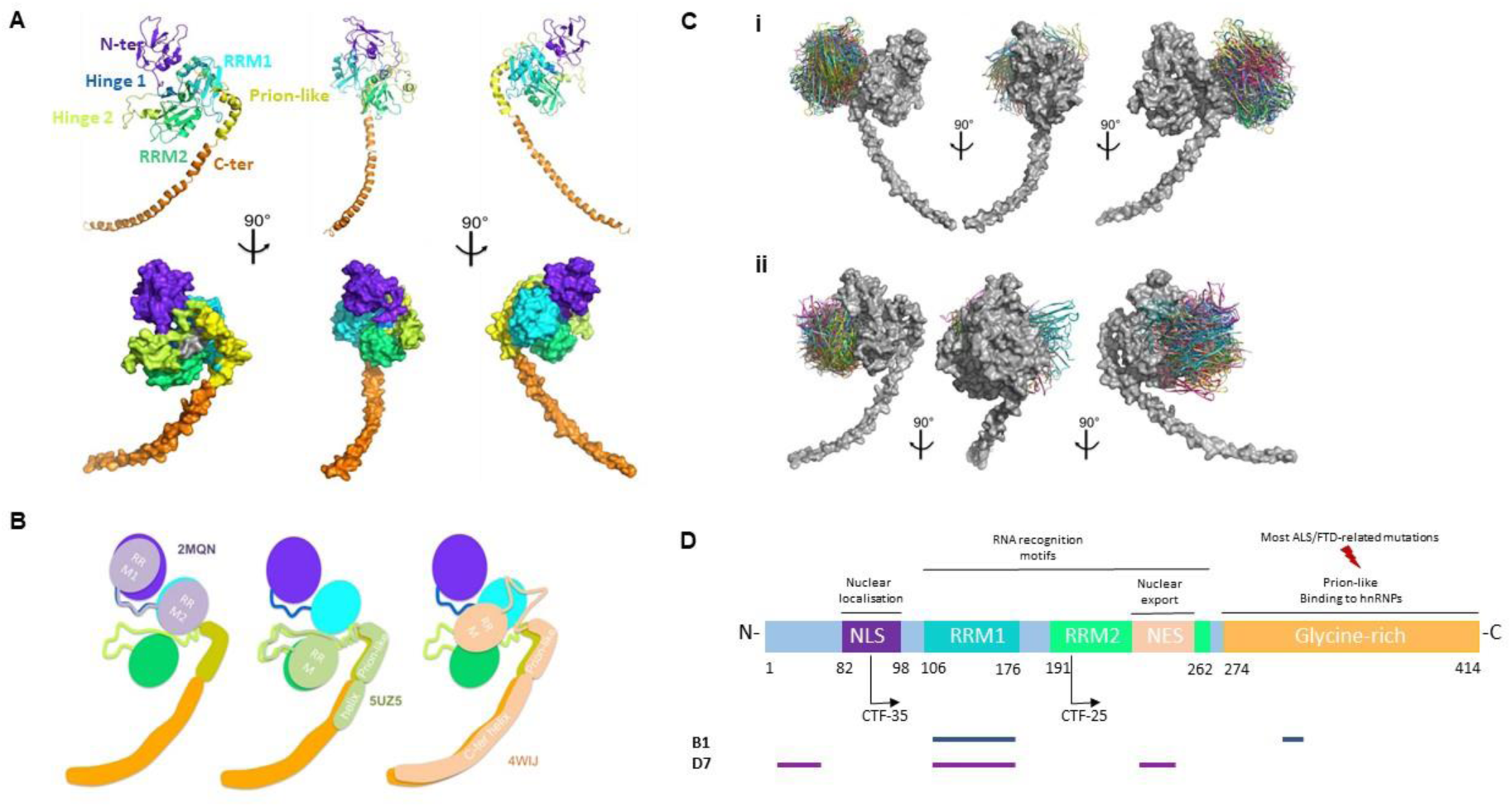
TDP-43 modeling and *in silico* epitope mapping. A) 3D model of full-length TDP-43. B) Templates used for modeling the relative orientations of the different domains of TDP-43. C) . View of the top 30 ranked predicted conformations for the complex between B1 and TDP-43 (i), and D7 and TDP-43 (ii). TDP-43 is shown in grey and the scFv in colors. D) *In silico* prediction of the interacting regions of TDP-43 with the scFvs B1 and D7.

The *in silico* binding site prediction required the 3D modeling of the scFv using templates for the V_H_ and V_L_ domains, with the orientation between the V_H_ and V_L_ domains being based on the V_H_ template orientation (data not shown). We then generated 30 ranked predicted conformations for the complex formed by the scFv with TDP-43 **(Fig.1C)**. From the top 30 ranked docking models, the residues of TDP-43 were scored for their probability of belonging to the epitope, and the residues of the scFv were scored for their probability of belonging to the paratope (data not shown). The scFv B1 could target a site containing the RRM1 domain and the prion-like domain **(Fig. 1D, blue line)**. As for the scFv D7, it was found to potentially bind a site consisting of the N-terminal domain, as well as the RRM1 and the RRM2 domains **(Fig. 1D, violet line)**.

### 3.3 scFvs B1 and D7 bound wtTDP-43

To verify whether the scFvs B1 and D7 interact with human wtTDP-43, we conducted an indirect ELISA on recombinant human full-length wtTDP-43 using purified B1, D7, and the control scFv, 13R4. The control wells containing increasing amounts of 13R4 showed no significant binding to TDP-43. However relative to the equivalent quantity of the control scFv, the wells incubated with increasing amounts of B1 and D7 exhibited a significant dose-dependent increase in absorbance **(Fig.2)**. Upon incubation with 5 µg/mL of B1 and D7, the signals increased 5-fold and 2-fold, respectively (B1, *p* = 0.004; D7, *p* = 0.03). Upon incubation with 10 µg/mL of B1 and D7, the signals increased 3-fold and 2-fold, respectively (B1, *p* = 0.004; D7, *p* = 0.005). In addition, surface plasmon resonance confirmed the binding of B1 and D7 to TDP-43, with a KD value in the 10^-8^ M range (data not shown). These results confirm that B1 and D7 physically interact with human wtTDP-43.

**Figure 2:**
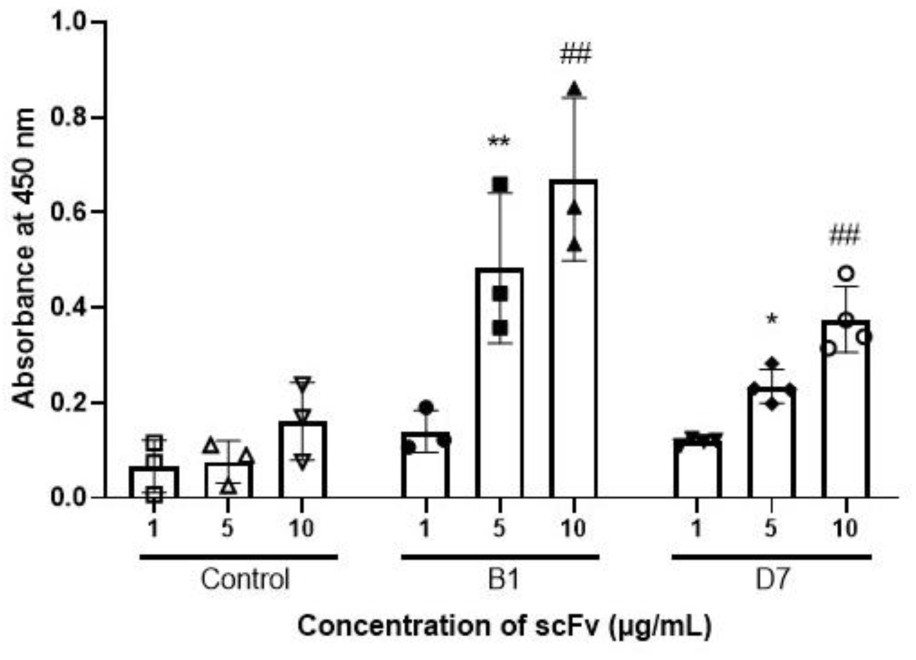
ELISA of purified scFv reveal the binding of B1 and D7 to recombinant wtTDP-43. Wells of 96-well plate coated with wtTDP-43 diluted to 1 µg/mL were incubated with the control scFv, B1, or D7 at increasing concentrations of 1 µg/mL, 5 µg/mL, and 10 µg/mL (x-axis). The control scFv did not show signs of binding to TDP-43 at any concentration, while B1 and D7 showed binding via increasing absorbances. Mann Whitney test: \*\**p*=0.004 or \**p*=0.03 versus 5 µg of control scFv; ##*p*<0.005 versus 10 µg of control scFv. N=6-7.

In cells not overexpressing TDP-43 and in which TDP-43 was mainly nuclear, the intrabodies seemed to be localized in the nucleus and the cytoplasm (**Fig.3A**). However, when TDP-43 was overexpressed and subsequently mislocalized to the cytoplasm (**Fig.4 A,B**), the localization of the intrabodies became mainly cytoplasmic, and colocalized with aggregated TDP-43 (**Fig3A**) MTT reduction assays revealed that the intrabodies had no significant toxicity on HEK293T cells **(Fig.3B)**. Similarly, the overexpression of TDP-43 with or without the selected intrabodies did not have an effect on cellular viability (**Fig.4C**). Interestingly, it was also noted that the turnover of the intrabodies of interest, but not the control, was much higher in the presence of pathological, overexpressed TDP-43 compared to control conditions **(Fig.S4)**.

**Figure 3:**
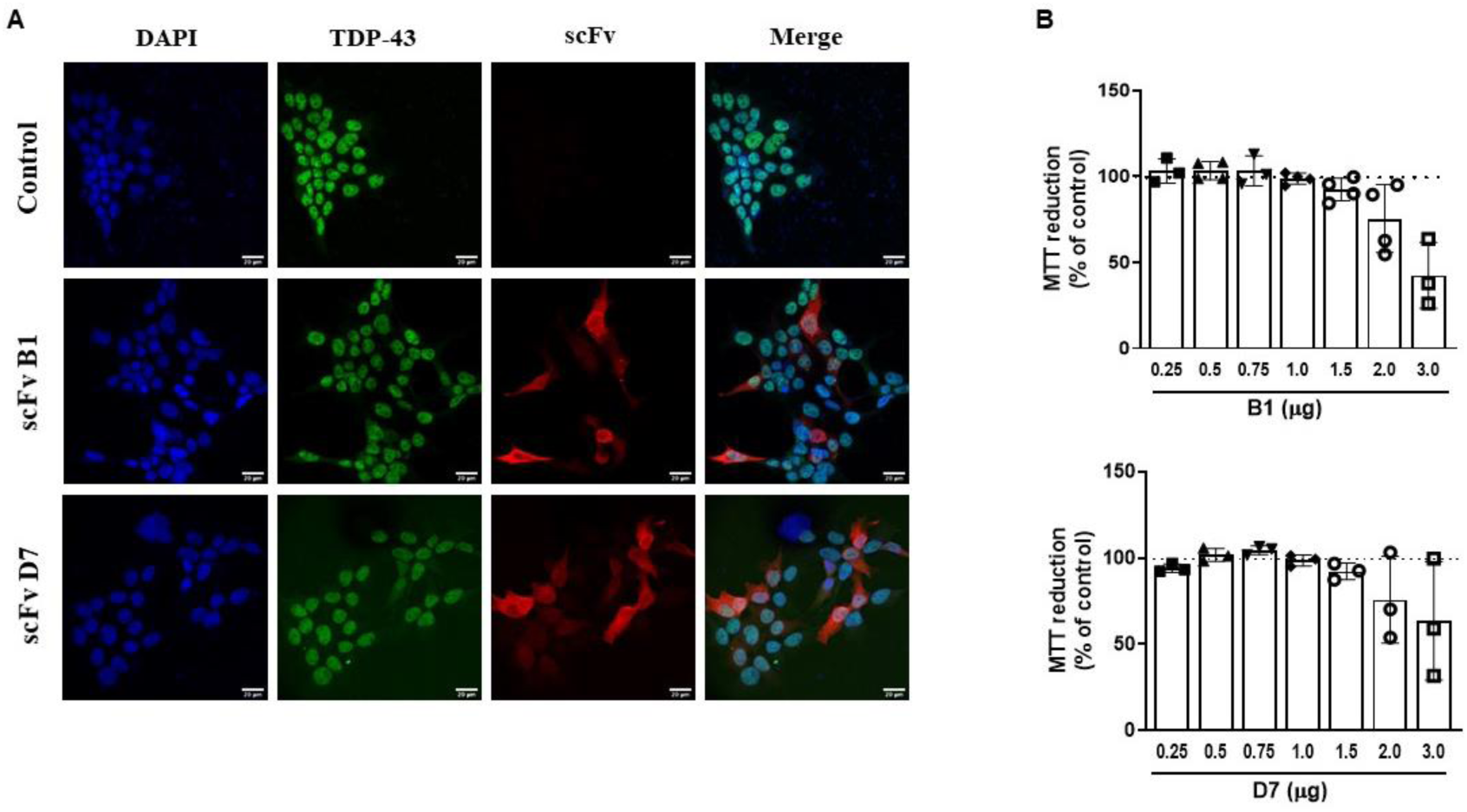
Expression and lack of cellular toxicity of the intrabodies. A) Immunofluorescence of HEK293T cells expressing the intrabodies. Scale bar = 20 µm (N=3). B) MTT reduction assay reveals the absence of toxicity of the intrabodies in HEK293T cells. Control: non-transfected cells. Quantity of scFv reflects the amount of plasmid transfected. One-way ANOVA, *p*<0.05, (N=3).

**Figure 4:**
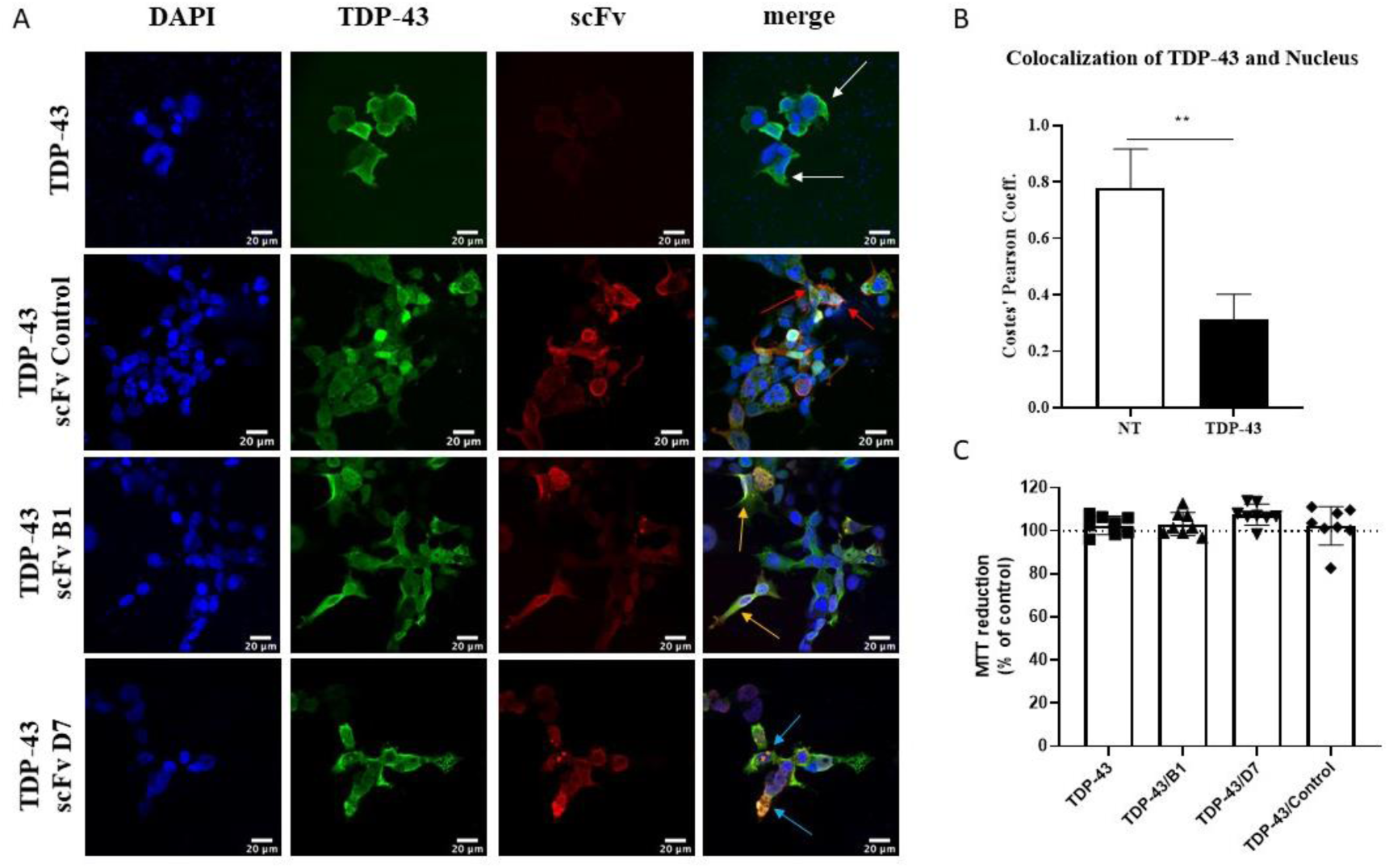
Intrabodies colocalize with mislocalized TDP-43 in the cytoplasm. A) Immunofluorescence of HEK293T cells co-transfected with TDP-43 (3 µg) and scFv (1.5 µg). White arrows: accumulation of TDP-43 in the cytoplasm. Red arrows: colocalization of control scFv with TDP-43. Orange arrows: colocalization of scFv B1 with TDP-43. Blue arrows: colocalization of scFv D7 with TDP-43. Scale bar = 20 µm (N=3). B) Colocalization of TDP-43 with the nucleus. Mann-Whitney test, \*\**p*=0.0016 (N=6). C) MTT reduction assay on HEK293T cells co-transfected with TDP-43 and scFv. Control = cells transfected with empty vector. Quantity of scFv reflects the amount of plasmid transfected. One-way ANOVA, *p*>0.5 (N=8).

### 3.4 D7 decreased the amount of the insoluble 35-kDa fragment

We quantified by Western blot the amount of full-length TDP-43 (TDP-43), the 35-kDa C-terminal fragment (TDP-35 CTF), and the 25-kDa C-terminal fragment (TDP-25 CTF) in the total lysate, soluble and insoluble fractions of HEK293T lysates. Upon transfection of cells with the plasmid overexpressing TDP-43, we observed a clear increase in the amounts of TDP-43 and TDP-35 CTF, as well as the appearance of TDP-25 CTF compared to the control (empty vector). The co-overexpression of the scFv D7 had no effect on the amount of either full-length TDP-43 nor its C-terminal fragments in the total or soluble protein fractions compared **(Fig.5)**. The intrabody B1 had no significant effect on the amount or solubility of TDP-43.

**Figure 5:**
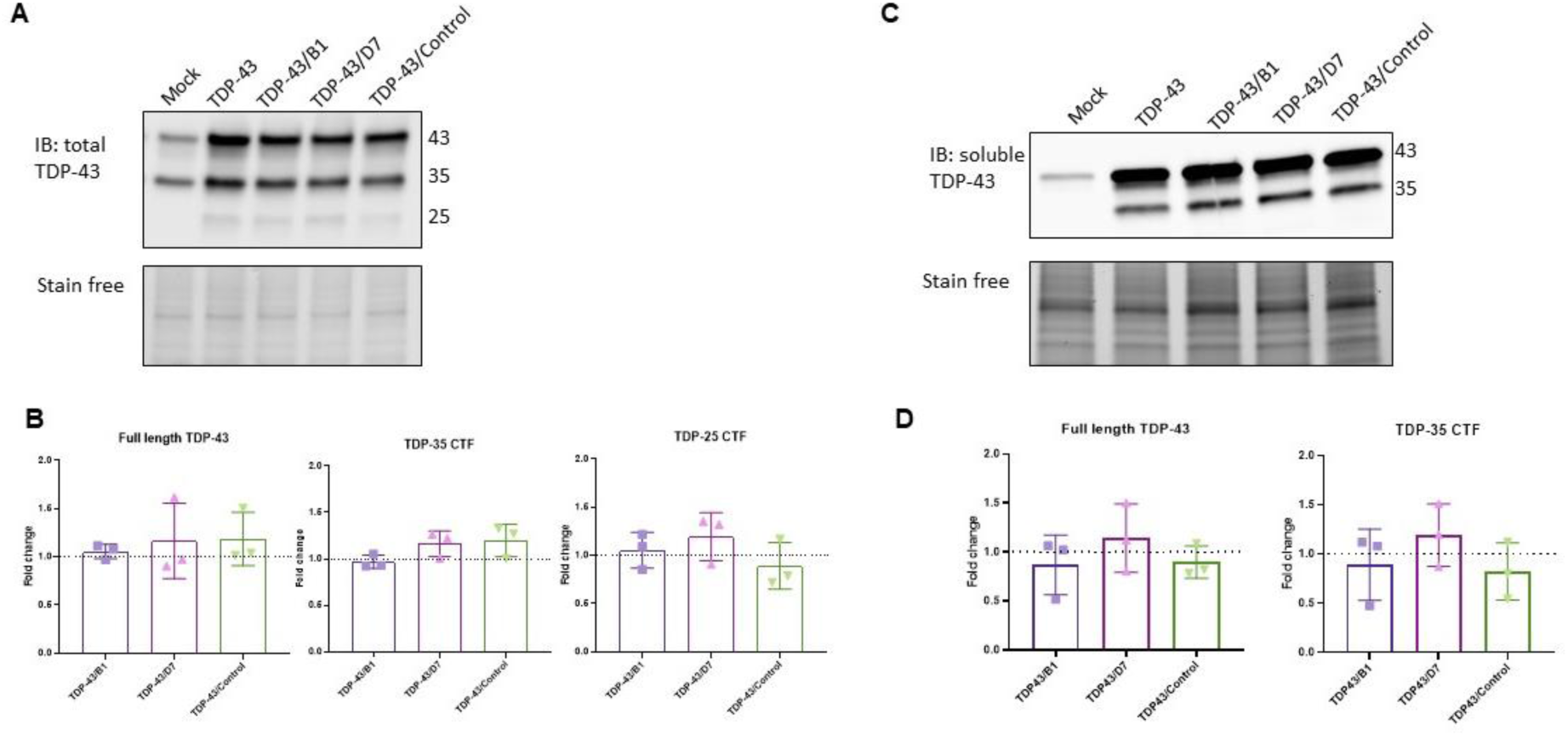
B1 and D7 have no significant effect on total and soluble TDP-43 or its CTFs. Immunoblots of TDP-43 in the total (A) and soluble (C) protein lysates of HEK293T cells. Quantification of TDP-43 forms in the total (B) and soluble (D) lysates normalized to the TDP-43 condition (dotted line). Mock: empty vector. TDP-43 condition: cells transfected with TDP-43-expressing plasmid and empty vector. TDP-43/B1, TDP-43/D7 and TDP-43/control: TDP-43-expressing plasmid and intrabody-expressing plasmid. Mann-Whitney test, *p*>0.5, (N=3).

In the insoluble protein fraction of HEK293T cells overexpressing TDP-43, we observed TDP-43 and TDP-35 CTF but not TDP-25 CTF **(Fig.6)**. The quantity of insoluble TDP-43 did not change upon overexpression of D7 (not shown), but the amount of insoluble TDP-35 CTF was significantly decreased (**Fig.6A,B**, *p* = 0.02). Interestingly, this effect of D7 was abolished when the cells were also treated with the proteasome inhibitor MG-132 (**Fig.6C,D**; *p* = 0.02). This effect was not observed with the autophagy inhibitor Bafilomycin A1 (*p* = 0.1). Concomitantly, when TDP-43 was overexpressed, the insoluble fraction of full-length TDP-43 and of TDP-35 CTF seemed to be mainly degraded by the proteasome, as evident by their significant increase when proteasomal degradation but not autophagy was inhibited (**Fig.6E,F**; *p* = 0.0286).

**Figure 6:**
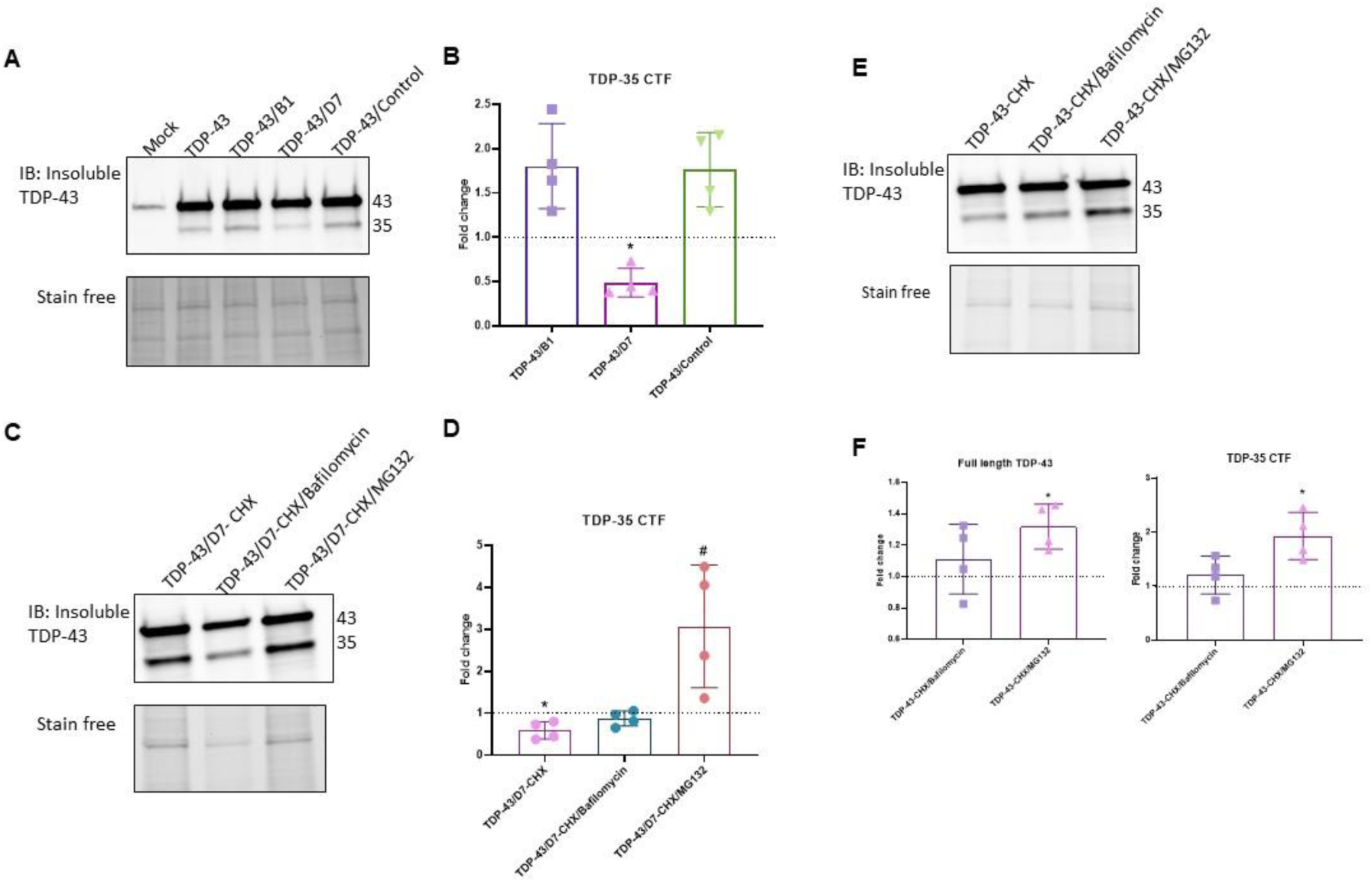
scFv D7 mediates the proteasomal degradation of the insoluble TDP-35 CTF. Immunoblots of TDP-43 in the insoluble protein fraction of HEK293T cells overexpressing TDP-43 and the intrabody in the absence (A) and presence (C) of autophagy and proteasome inhibitors. Quantification of the insoluble TDP-35 CTF normalized to the TDP43 condition (dotted line) in the absence (B) and presence (D) of autophagy and proteasome inhibitors. CHX: cycloheximide (10 µg/mL), Bafilomycin A1 (300 nM), MG-132 (0.5 µM). \**p*=0.0286 vs TDP43; *#p=*0.0286 vs TDP43/D7; Mann-Whitney test (N=4). E) Immunoblot of TDP-43 in the insoluble protein fraction of HEK293T cells overexpressing TDP-43 and treated with autophagy or proteasome inhibitors. F) Quantification of insoluble TDP-43 forms normalized to the vehicle condition (dotted line). \**p*=0.0286, Mann-Whitney test (N=4).

### 3.5 The metabolomic profile of cells was affected by TDP-43 and scFv expression

We analyzed the metabolomic profiles of HEK293T transfected either with TDP-43 plasmid alone or co-transfected with TDP-43 plasmid and each intrabody plasmid (D7, control 13R4) to evaluate (1) the metabolic effect of scFv administration, (2) metabolic modification associated with TDP-43 overexpression, and (3) correction of TDP-43–linked metabolic changes by D7. We identified a total of 220 metabolites in HEK293T cells. Lists of the 10% of metabolites that were the most upregulated and downregulated are presented in **Tables S2 and S3**, respectively. Compared with cells transfected with the empty vector, the majority of the metabolites impacted seem to be common to the two intrabodies **(Fig. 7A)**. They represent the global effect of the overexpression of an scFv on naïve cells, with only a few metabolites specific to D7. The same was observed when comparing the conditions of TDP-43 overexpression and TDP-43/scFv co-transfection, with a majority of metabolites being common to the two intrabodies **(Fig.7B)**. The metabolic effects of D7 also seemed to vary depending on whether TDP-43 was overexpressed or not **(Fig.7C)**, suggesting different effects when TDP-43 is pathological. We were able to identify three metabolites (L-palmitoylcarnitine, PC[36:4], 4-hydroxy-phenylglycine) that were altered when TDP-43 was overexpressed and that returned to baseline levels (empty vector condition) when the scFv D7 was co-expressed **(Fig.7D)**.

**Figure 7:**
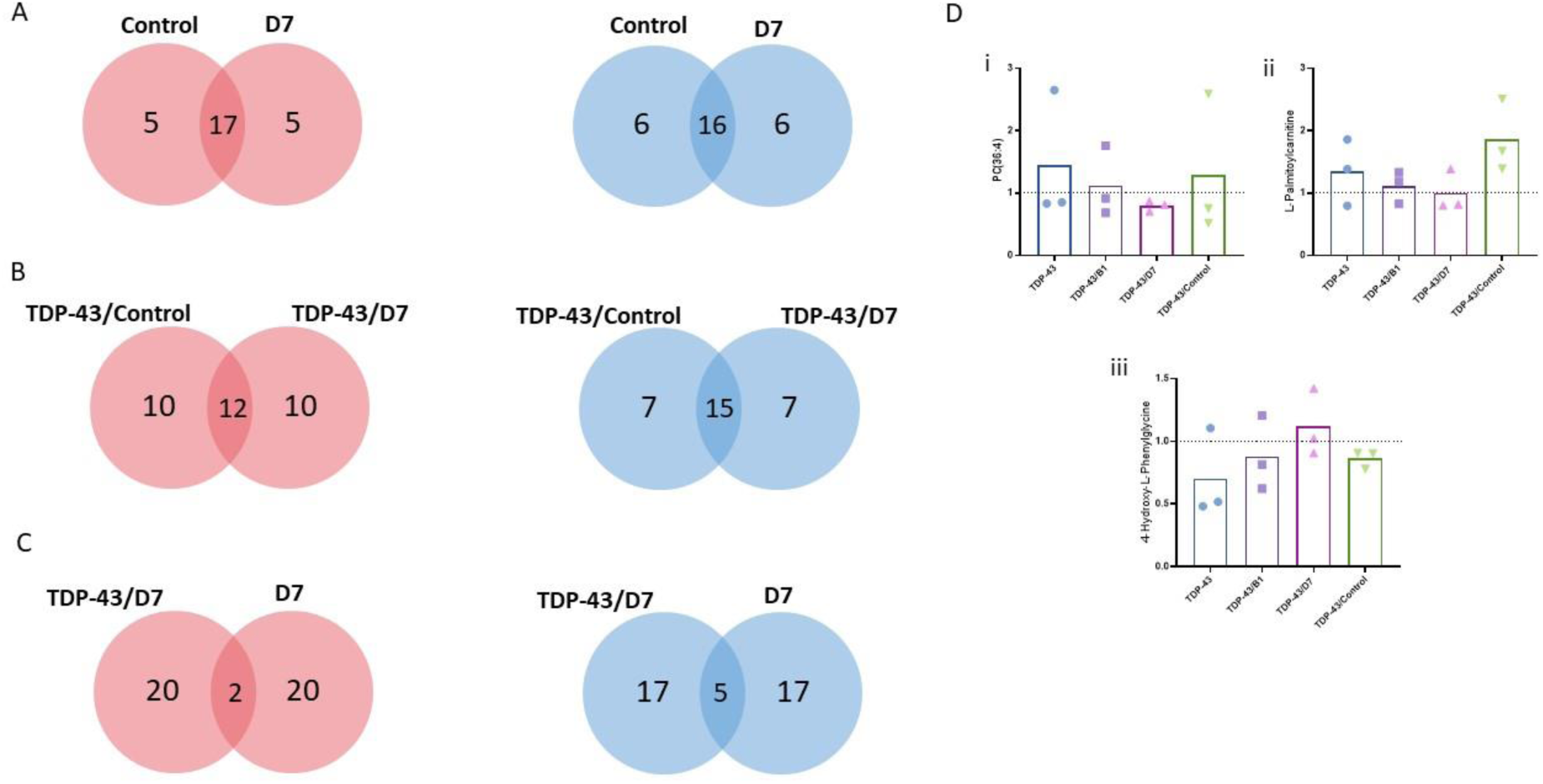
Metabolomic profiles of HEK293T cells. Venn diagram representing the number of specific and common metabolites that increased (red) or decreased (blue) within each condition upon A) overexpression of the scFv (values normalized to the mock condition), (B) overexpression of TDP-43 and the scFv together (values normalized to the TDP-43 condition), C) transfection with the scFv with or without TDP-43 overexpression (values normalized to the TDP-43 condition). D) Quantification of some of the metabolites of interest. i) PC(36:4), ii) L-palmitoylcarnitine, iii) 4-hydroxy-L-Phenylglycine. Values were normalized to the vehicle condition (dotted line).

## 4. Discussion

TDP-43 plays a crucial role in maintaining cellular homeostasis by regulating gene expression, RNA processing, and RNA stability(2). In disease states, the protein is found mislocalized to the cytoplasm where it undergoes fragmentation and post-translational modifications and aggregates. Targeting TDP-43 proteinopathy has been the focus of many research groups given the important role that this protein plays in many neurodegenerative diseases (14,38).

scFv intrabodies can offer a highly specific approach to directly target intracellular antigens and have gained growing interest as a therapeutic approach for neurodegenerative diseases(39). Two previous studies have selected scFvs for their specific binding to the RRM1 and RRM2 domains of TDP-43(40,41). Similarly, another team used *in silico* predictions to design nanobodies that intracellularly target the RRM domains of TDP-43(42). In 2020, a study used phage display to select diagnostic scFv clones that can recognize TDP-43 variants found in FTD(43). These studies highlight how targeting TDP-43 can pave the way for developing broad-spectrum therapies that might be effective across various conditions, but can also help in the diagnosis of TDP-43 proteinopathies.

We decided to follow an untargeted high-throughput screening approach using a synthetic scFv phage display library. This technique allows the screening of big libraries that are not biased towards a specific antigen, in order to identify scFvs against different potential sites on TDP-43. In addition, we based our screening on wtTDP-43, which is the form most commonly implicated in TDP-43 proteinopathies(2). We were able to identify four different TDP-43-specific scFv clones, two of which (B1 and D7) were retained for subsequent experiments. By not engineering the scFvs against a specific domain of TDP-43, we were able to generate two clones that targets different sites of TDP-43 spanning various 3D folded domains.

After verifying that the scFvs D7 and B1 specifically bind to wtTDP-43, we expressed them intracellularly as intrabodies in a HEK293T cell model of TDP-43 proteinopathy. A main challenge of scFvs targeting TDP-43 is ensuring that they do not target functional nuclear TDP-43. In our case, the intrabodies were found to be mainly cytoplasmic in the presence of mislocalized TDP-43. The intrabodies B1 and D7 also colocalized with aggregated TDP-43, which was not the case for the control intrabody. We also detected no effect of our intrabodies on soluble TDP-43 by western blot.

Given that overexpressing TDP-43 for 48 h in HEK293T cells did not lead to significant cell death compared to the control, we could not use cellular viability as a main criterion to assess the effect of the intrabodies on TDP-43-induced cellular toxicity. We confirmed however that our intrabodies did not negatively affect the viability of cells that were already under TDP-43-induced stress. Given the progressive and complex nature of ALS, and the interplay between several malfunctioning mechanisms, it would be interesting to assess more subtle cellular and metabolic changes to evaluate the effectiveness of some TDP-43-targeting approaches. In our case where TDP-43 and the intrabodies are co-overexpressed at the same time and only for 48 h, we can assume that the effect of the scFv occurs on pathological changes that appear at the earliest stages of the disease.

TDP-43 aggregation was represented by the amount of protein recovered in the detergent-insoluble fraction of cell lysates. One study has previously demonstrated that visible TDP-43 inclusions under confocal microscopy were not necessarily insoluble(44); based on this data we used western blot analyses to quantify TDP-43 insolubility. Overexpression of TDP-43 led to the fragmentation of the protein, with the appearance of TDP-35 and TDP-25 CTF. Interestingly, the overexpression of the intrabody D7 led to a decrease in the amount of insoluble TDP-35 CTF in HEK293T cells while the total amount of TDP-43 remained unchanged. The intrabody B1 did not have any effect on the amount or insolubility of TDP-43. The effect of D7 on the insoluble TDP-35 CTF level was reversed upon inhibition of the proteasome, but not upon inhibition of autophagy. This is in line with our observation that upon overexpression of TDP-43 alone, the detergent-insoluble fraction of TDP-43 and TDP-35 CTF was mainly degraded by the proteasome. Together, these results suggest that D7 enhances the proteasomal degradation of the detergent-insoluble TDP-35 CTF while sparing functional TDP-43. It remains unclear to us why D7 targeted TDP-35 CTF instead of TDP-43, but we speculate that it might be due to different conformations of TDP-43 and TDP-35 inside cells making the epitopes on TDP-35 more accessible to D7. For conditions like TDP-43 proteinopathy, promoting the degradation of toxic aggregation-prone proteins can alleviate cellular stress.

CTFs of TDP-43 have been found to be associated with several neurological disorders such ALS(3), FTD(45), Alzheimer’s disease, corticobasal degeneration(46), Pick’s disease(47), Parkinson’s disease(48), and traumatic brain injury(49). TDP-35 CTF is a product of the proteolytic cleavage of TDP-43 at the level of its N-terminus(9,50). Given that TDP-35 CTF includes the RNA-binding domains of TDP-43, particularly the RRM1 domain, this fragment retains its ability to associate strongly with different RNA species(51). TDP-35 lacks the nuclear localization signal of TDP-43, and predominately localize in the cytoplasm and form stress granules or aggregates that negatively affect RNA processing (52). One study showed that TDP-35 CTF can redirect TDP-43 from its normal nuclear location to cytoplasmic aggregates, and that this process depends crucially on RNA binding(53,54). In addition, TDP-35 was found to be even more aggregation-prone than TDP-43 or TDP-25, and to seed to aggregation of TDP-43(54). All these studies highlight the importance of targeting TDP-35 CTF as a strategy to counteract or halt the progression of TDP-43 proteinopathy.

The dysfunction of TDP-43 in neurodegenerative diseases is known to cause significant cellular alterations including disturbances in the metabolipidomics profile(11,55,56). D7 exhibited the ability to restore the levels of different metabolites altered by TDP-43 overexpression. In HEK293T cells, L-palmitoylcarnitine and glycerophospholipid PC(36:4) were decreased and restored back to basal levels by D7. Interestingly, our group previously reported perturbations in the levels of numerous carnitines and glycerophospholipids in HEK293T cells overexpressing TDP-43(26). L-palmitoylcarnitine is a 23-carbon acylcarnitine that is increased is ALS(57,58), suggesting an impaired beta-oxidation. Phosphatidylcholine PC(36:4) was also one of the main molecules that increased in ALS patients compared to controls, and was found to correlate with disease progression(59,60). D7 also seemed to increase back to normal the level of 4-hydroxy-phenylglycine, a known inhibitor of carnitine palmitoyltransferase-1 (CPT1)(61). Various studies have pinpointed alterations in CPT1 in TDP-43 proteinopathies(57,62). For instance, downregulation of CPT1 was found to alleviate pathological alterations and symptoms in a mouse model of ALS(63). Metabolomic analysis seems to point towards a clear involvement of lipid metabolism in TDP-43 proteinopathy which was targeted by D7.

## 5. Conclusions

Using phage-display, we identified two main scFv clones that can bind to human wtTDP-43. Of the identified scFvs, intrabody D7 was able to reduce the toxic burden of TDP-43 in cells, particularly by decreasing the levels of insoluble TDP-35 CTF, and alleviate some metabolic alterations linked to TDP-43 proteinopathy. This can serve as tool to help decipher the role of TDP-35 CTF in health and disease and the importance of targeting cellular alterations linked to TDP-43 proteinopathy such as metabolomic changes.

## Acknowledgements

The authors would like to thank Professor Giles Lalmanach for helpful discussions about the ELISA test; Dr Eric Reiter and Dr Anne Poupon for the MAbSilico analyses; and Professor Hervé Watier and all the LabEx network for their support. This publication has been funded with support from Région Centre Val de Loire and the French National Research Agency under the program “Investissements d’avenir” Grant Agreement LabEx MAbImprove: ANR-10-LABX-53. We would like to thank the ARSLA for financial support. Funding sources had no participation in study design; in the collection, analysis, and interpretation of data; in the writing of the report; and in the decision to submit the article for publication.

## Author Contributions

**Y.A.O., R.H.**: Conceptualization, data curation, formal analysis, investigation, methodology, visualization, writing (original draft); **H.A.**: formal analysis, methodology, visualization, writing (original draft). **A.C, J.A., A.D., S.H., J.B. and A.L**.: investigation; **A.D., S.O., P.E., P.V., C.R.A., P.C. and O.H.**: writing (review & editing); **P.M.**: Supervision, resources, writing (review & editing); **D.L.**: conceptualization, methodology, investigation, validation, supervision, writing (review & editing); **H.B.**: supervision, project administration, funding acquisition, writing (review & editing). All authors read and approved the final manuscript.

