## Supplementary data for "scFv intrabody targeting wildtype TDP-43 presents protective effects in a cellular model of TDP-43 proteinopathy"

#### Supplementary figures

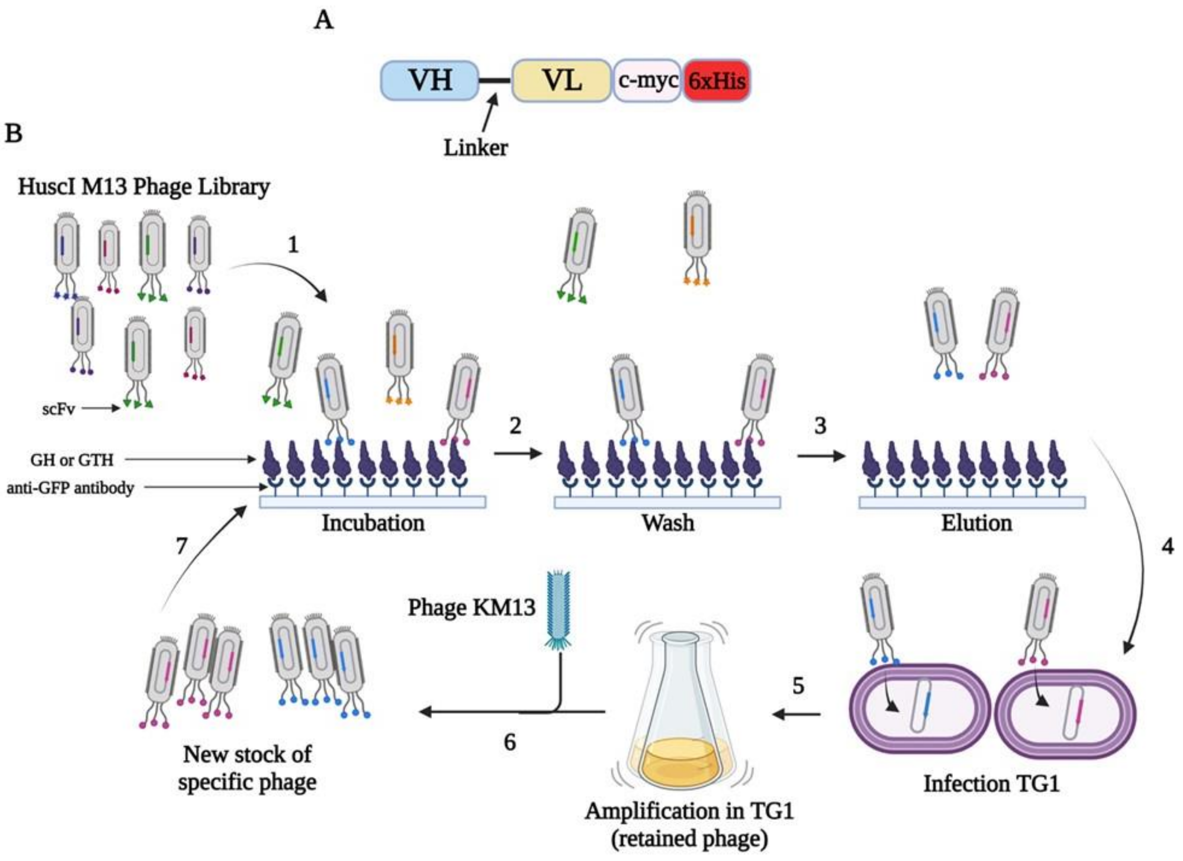


**Figure S1: Flow chart illustrating the phage display screening strategy of scFv molecules.** A) General structure of scFv protein. Once expressed as protein isolated from phage, it contains C-terminal c-myc and 6xHis tags. B) The HuscI library contained M13 phage that expressed copies of one scFv at their tail. The library that was depleted against anti-GFP antibody and against GFP-6xHis (GH) was incubated with GFP-wtTDP-43-6xHis (GTH) immobilized on anti-GFP antibody (1). After washing (2), the adhered phage were eluted by trypsin (3) and induced to infect TG1 (4). Following amplification of infected TG1 (5), new phage particles were produced (6) containing amplified specific phage. These phage were re-incubated with GTH (7). The cycle was repeated a total of 5 times. Figure made with Biorender.com.

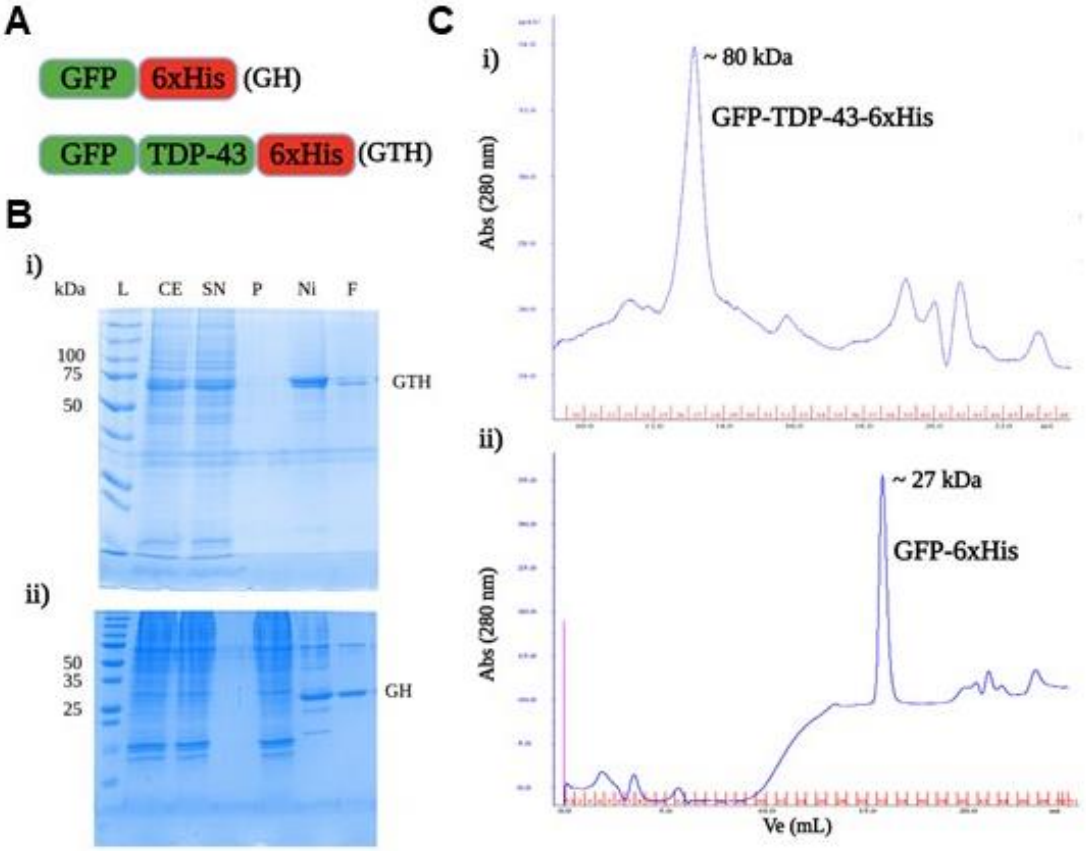


**Figure S2: Purification of recombinant antigens for phage display.** A) Structures of the recombinant proteins GFP-6xHis (GH) and GFP-TDP-43-6xHis (GTH) of 28 kDa and 75 kDa, respectively. B) SDS-PAGE gel stained by Coomassie reagent representing the purification of i) GTH and ii) GH. L: ladder; CE: crude extract of lysate. SN: supernatant of lysate. P: pellet of lysate. Ni: pooled eluted protein from affinity chromatography. F: final product following dialysis. C) Gel filtration profile of dialyzed i) GTH and ii) GH. The chromatogram reflects highly pure monomers of each protein. Ve: elution volume.

A

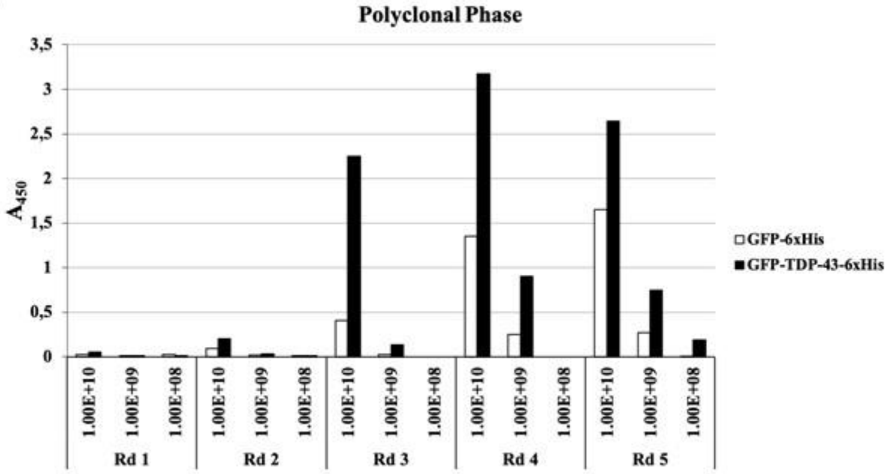
Polyclonal Phase
$A_{450}$
3,5
3
2,5
2
1,5
1
0,5
0
1.00E+10
1.00E+09
1.00E+08
Rd 1
Rd 2
Rd 3
Rd 4
Rd 5
GFP-6xHis
GFP-TDP-43-6xHis

B

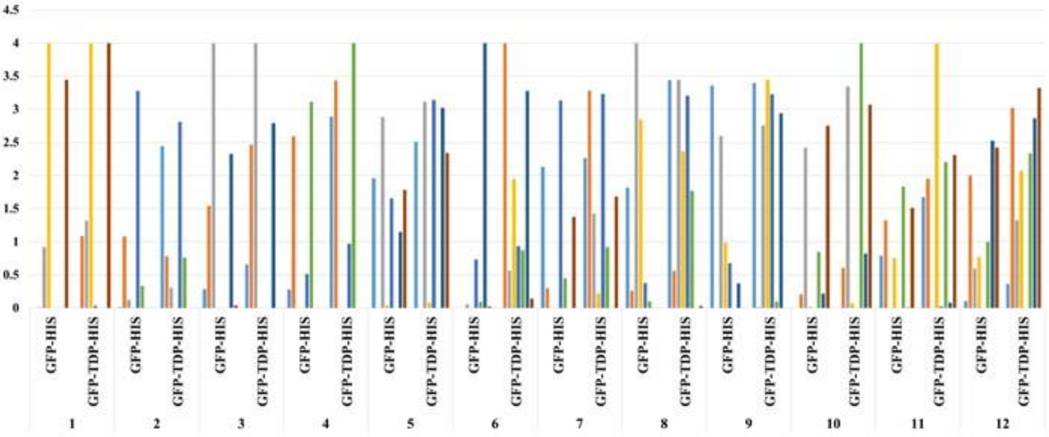
A 450 nm
4.5
4
3.5
3
2.5
2
1.5
1
0.5
0
GFP-HIS
GFP-TDP-HIS
1
2
3
4
5
6
7
8
9
10
11
12

C

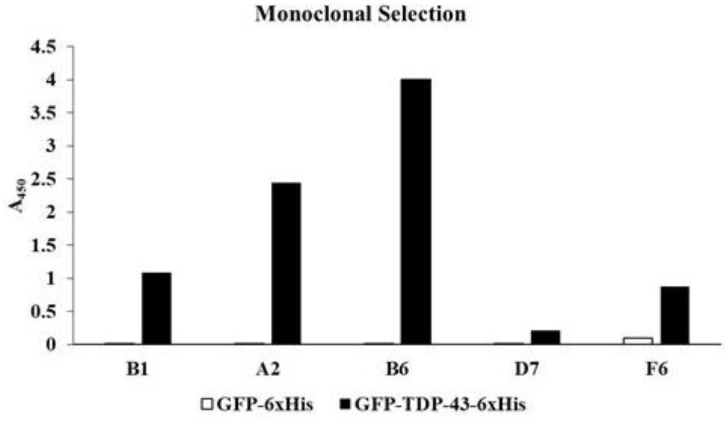
Monoclonal Selection
$A_{450}$
4.5
4
3.5
3
2.5
2
1.5
1
0.5
0
B1
A2
B6
D7
F6
GFP-6xHis
GFP-TDP-43-6xHis

**Figure S3:** A) Amplification of polyclonal phage against GTH. Each round of eluted phage was tested by ELISA on wells coated with GH and GTH antigen. The x-axis represents the 3 dilutions in phage/mL used on coated wells. The white bar represents the absorbance at 450 nm for GH incubation, while the black bar represents the absorbance for GTH incubation. Round 4 appears to contain the most phage with affinity for GTH. B) Monoclonal screen of selected phage from round 4 for binding to GH and GTH. C) Identification of monoclonal anti-TDP-43 scFv-phage display from TG1. Phage with distinct scFv sequences from wells B1, A2, B6, and D7 were identified as anti-TDP-43 because no absorbance was detected with GH. Phage from well F6 is represented as an example of non-specific binding because absorbance was also detected when incubated with the GH control.

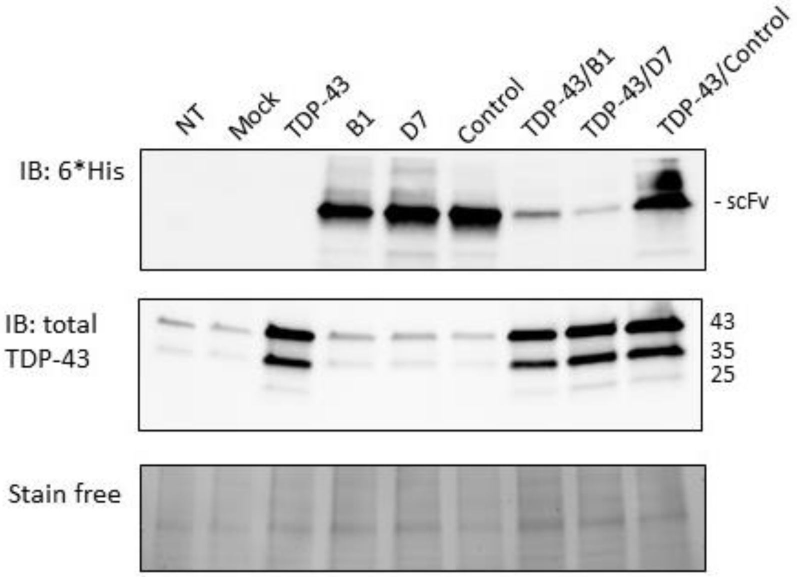


**Figure S4: Overexpression of 6*His scFv and TDP-43 in HEK293T cells.** Immunoblots showing the expression of the intrabody when expressed alone and when co-overexpressed with TDP-43. NT: non-transfected. Mock: cells transfected with the empty vector. TDP-43: cells transfected with TDP-43-expressing plasmid and empty vector. B1, D7, control: cells transfected with intrabody-expressing plasmid and empty vector. TDP-43/B1, TDP-43/D7 and

TDP-43/control: cells transfected with TDP-43-expressing plasmid and intrabody-expressing plasmid.

### Supplementary tables

#### Table S1: List of antibodies used in this study

| Name of the antibody | Manufacturer (catalog #) | Species raised in | Dilution |
|---|---|---|---|
| HRP-conjugated anti-6*His | Proteintech (HRP-66005) | Mouse monoclonal | 1:100 000 (WB) 1:10 000 (ELISA) |
| Anti-6*His | Proteintech (HRP-66005-1-Ig) | Mouse monoclonal | 1:1000 (IF) |
| Anti-c-Myc | Thermo Fisher Scientific (13-2500) | Mouse monoclonal | 1:2000 (ELISA) |
| HRP-conjugated M13 bacteriophage | Sino Biological ( 11973-MM05T-H) | Mouse monoclonal | 1:2000 (ELISA) |
| Anti-TDP-43 (C-terminal) | Proteintech (12892-1-AP) | Rabbit polyclonal | 1 :1000 (IF) 1 :5000 (WB) |
| Anti-NFκB Antibody, p65 subunit | Sigma Aldrich (MAB3026) | Mouse monoclonal | 1:1000 (WB) |
| Alexa Fluor™ 488-conjugated anti-rabbit | Thermo Fisher Scientific (A-11034) | | 1 :500 (IF) |
| Alexa Fluor™ 594-conjugated anti-mouse | Thermo Fisher Scientific (A-11005) | | 1:500 (IF) |
| HRP-conjugated anti-rabbit | Promega (W401B) | | 1:10 000 (WB) |
| HRP-conjugated anti-mouse | Promega (W402B) | | 1:10 000 (WB) |

#### Table S2: List of most upregulated metabolites within each condition in HEK293T cells

| TDP-43 vs Empty | 13R4 vs Empty | D7 vs Empty | TDP-43/13R4 vs TDP-43 | TDP-43/D7 vs TDP-43 |
|---|---|---|---|---|
| Creatine phosphate | Deoxyadenosine monophosphate | Deoxyadenosine monophosphate | Cysteic acid | Cysteic acid |
| Citicoline | Inosine | 7-dehydrocholesterol | Ornithine | Diethanolamine |
| PC(36:4) | N-alpha-acetyl-lysine | L-oleoyl-rac-glycerol | Urocanic acid | Glucose-1-phosphate |
| L-glutamine | C14-carnitine | Inosine | L-histidinol | L-histidinol |
| Glycerophosphocholine | Galactitol | LysoPE(20:4) | C14-carnitine | Urocanic acid |
| L-palmitoylcarnitine | L-histidinol | Galactitol | Homogentisate | Trigonelline |
| Cortisone | Elaidic acid | Ornithine | 3-methylglutaric acid | PC(35:4) |
| C14-carnitine | 7-dehydrocholesterol | LysoPE(18:0) | Diethanolamine | Spermidine |
| Aminoadipic acid | L-oleoyl-rac-glycerol | Elaidic acid | Spermidine | Methyl jasmonate |
| Sarcosine | LysoPE(20:4) | Thymidine | PC(36:5) | 4-hydroxy-L-phenylglycine |
| 7-dehydrocholesterol | Thymidine | SM(32:2) | L-serine | N-acetylmannosamine |
| Urocanic acid | Pterin | LysoPE(16:0) | Succinic acid | 3-methylglutaric acid |
| Creatinine | Ornithine | N-acetylneuraminate | PC(35:4) | Spermine |
| L-proline | N-acetylneuraminate | PE(16:0/18:2) | Spermine | D-galactosamine |
| Taurine | 4-hydroxy-L-phenylglycine | N-acetyl-L-methionine | Glucose-1-phosphate | LysoPC(20:4) |
| LysoPE(20:4) | PC(36:4) | PC(36:4) | Glyceraldehyde-3-phosphate diethyl acetal | Tyramine |
| C18-carnitine | SM(32:2) | Pterin | PC(37:4) | PC(37:5) |
| Oxoglutaric acid | LysoPE(18:0) | PAF-C16 | Methyl jasmonate | 10-hydroxy capric acid |
| L-aspartic acid | Glucose-1-phosphate | PC(40:9) | L-palmitoyl-carnitine | Omega-hydroxydodecanoic acid |
| N-aceyl-L-phenylalanine | LysoPC(14:0) | N-alpha-acetyl-lysine | C18-carnitine | Sorbate |

| | | | | |
|---|---|---|---|---|
| Inosine | Xanthine | LysoPC(16:0) | Trigonelline | All-trans-retinoic acid |
| Alpha-glucose | N-acetyl-L-methionine | 4-hydroxy-L-phenylglycine | LysoPC(20:4) | LysoPE(20:4) |

#### Table S3: List of most downregulated metabolites within each condition in HEK293T cells

| **TDP-43 vs Empty** | **13R4 vs Empty** | **D7 vs Empty** | **TDP-43/13R4 vs TDP-43** | **TDP-43/D7 vs TDP-43** |
|---|---|---|---|---|
| Glucose-1-phosphate | Aminoadipic acid | Phosphorylcholine | Inosine-5'monophosphate | Inosine |
| Spermidine | Uridine-5'-diphosphoglucose | Citicoline | 7-dehydroxycholesterol | Inosine-5'monophosphate |
| Deoxyadenosine monophosphate | Citicoline | Aminoadipic acid | PE(22:6/18:1) | PE(16:0/22:6) |
| Trigonelline | Spermidine | Uridine-5'-diphosphoglucose | Inosine | PC(40:9) |
| Spermine | Creatine phosphate | Spermidine | Cytidine | Uracil |
| 4-hydroxy-L-phenylglycine | Phosphorylcholine | Creatine phosphate | Fumarate | Cytidine |
| PC(40:8) | L-glutamine | L-glutamine | Lactate | Pyridoxal-5'-phosphate |
| Sphinganine | Glutathione | Oxoglutaric acid | Glutathione | Uridine |
| Uridine-5'-monophosphate | O-succinyl-L-homoserine | Trigonelline | Pyridoxal-5'-phosphate | Lactate |
| Nicotinic acid | Propionylcarnitine | Creatinine | Uracil | PC(36:4) |
| Methyl jasmonate | S-adenosylmethionine | Propionylcarnitine | Malate | L-acetylcarnitine |
| Ornithine | L-acetylcarnitine | L-acetylcarnitine | PC(40:9) | 7-dehydroxycholesterol |

| | | | | |
|---|---|---|---|---|
| PC(37:4) | Creatinine | PC(o-20:4/2:0) | Uridine | PC(31:0) |
| PC(40:1) | Thiamine | SM(38:1) | Glucosamine-6-phosphate | Citicoline |
| Glyceraldehyde-3-phosphate diethyl acetal | Oxoglutaric acid | PC(o-20:6/2:0) | L-proline | L-palmitoylcarnitine |
| SM(38:2) | 3-dehydroxycarnitine | Succinic acid | L-arabitol | Fumarate |
| Riboflavin | Allantoin | Butyrylcarnitine | L-acetylcarnitine | Hypoxanthine |
| PC(39:4) | Glutamate | 3-dehydroxycarnitine | PE(16:0/22:6) | Glycerophosphocholine |
| L-serine | L-aspartic acid | Glutathione | N-acetyl-L-glutamic acid | Uridine-5'-diphosphoglucose |
| PC(38:8) | PC(35:4) | Sarcosine | Glycerophosphocholine | N-acetyl-L-methionine |
| All-trans-retinoic acid | Creatine | PC(35:4) | Uridine-5'-diphosphoglucose | Succinic acid |
| Elaidic acid | Sarcosine | Thiamine | L-methionine | Glutathione |
